# Development and Validation of an MRI-Derived Head-Neck Finite Element Model

**DOI:** 10.1101/2023.02.12.528203

**Authors:** Hossein Bahreinizad, Suman K. Chowdhury, Gustavo Paulon, Leonardo Wei, Felipe Z. Santos

## Abstract

**Purpose:** This study aimed to develop and validate a magnetic resonance imaging (MRI)-based biofidelic head-neck finite element (FE) model comprised of scalp, skull, CSF, brain, dura mater, pia mater, cervical vertebrae, and discs, 14 ligaments, and 42 neck muscles.

**Methods:** We developed this model using head and neck MRI images of a healthy male participant and by implementing a novel meshing algorithm to create finer hexahedral mesh structures of the brain. The model was validated by replicating four experimental studies: NBDL’s high acceleration profile, Ito’s frontal impact cervical vertebrae study, Alshareef’s brain sonomicrometry study, and Nahum’s impact study.

**Results:** The results showed reasonable geometrical fidelity. Our simulated brain displacement and cervical disc strain results were close to their experimental counterparts. The intracranial pressure and brain stress data of our head-only model (excluding neck structures and constraining the base of the skull) were similar to Nahum’s reported results. As neck structures were not considered in Nahum’s study, the FE results of our head-neck model showed slight discrepancies. Notably, the addition of neck structures (head-neck model) reduced brain stress values and uncovered the brain’s intracranial pressure dynamics, which the head-only model failed to capture. Nevertheless, the FE simulation results showed a good agreement (r > 0.97) between the kinematic responses of the head-neck model and NBDL’s experimental results.

**Conclusion:** The developed head-neck model can accurately replicate the experimental results and has the potential to be used as an efficient computational tool for brain and head injury biomechanics research.

**Statements and Declarations:** This work was primarily supported by the U.S. Department of Homeland Security (70RSAT21CB0000023). The MRI data acquisition was supported by the Texas Tech Neuroimaging Center.

## Introduction

In the field of biomechanics, computational head-neck models based on finite element (FE) methods play a pivotal role in assessing the responses of the brain and other head-neck components to various impact scenarios. These scenarios encompass a wide range, from contact sports [1] and motor vehicle accidents [2] to battlefield scenarios [3] and occupational settings [4]. FE modeling proves invaluable in understanding the spatial and temporal distribution of stress waves following head impacts and evaluating whether they exceed the strength and deformation tolerance limits of the constituent structures. This capability is especially crucial, given the challenge of measuring such responses through in-vivo experiments. While recent advancements in computational power and the availability of tissue material properties have made modeling the complex geometries of head-neck structures more accessible than ever, there remains a significant gap in the development of sophisticated and biologically accurate head-neck FE models.

The computational modeling of head-neck structures remains challenging due partially to the large variability in their mechanical properties. For instance, scalp exhibits high damping characteristics and linear elastic behavior under mechanical loads [5]. Skull shows a linear viscoelastic behavior with a higher stiffness in the elastic region [6]. The brain was found to display a distinctive non-linear viscoelastic behavior [7]. Moreover, research has demonstrated that white matter exhibits a notably firmer mechanical response in contrast to grey matter [8], underscoring the critical need for separate modeling of these regions. In addition, modeling the fluid behavior of cerebrospinal fluid (CSF) is challenging and requires CSF-skull and CSF-brain interfaces to be defined as complex fluid-structure interfaces. However, the interactions between most head-neck substructures, including CSF-brain and CSF-skull interfaces, were modeled as tied and/or sliding contact interfaces in order to reduce additional complexities in the FE modeling and simulation process [9, 10]. Besides material properties, the accuracy in geometry generation and the choice of FE meshing method (mesh type, element size, and mesh quality) are crucial to ensure that the head-neck models provide accurate numerical solutions [11]. This becomes even more critical when developing a detailed head-neck model because coupling many inaccurate surfaces in a complex model leads to singularities and divergence [11]. Similarly, an inappropriate meshing of a biological structure, even for geometrically accurate structures, was found to yield misleading numerical results [11]. For example, some previous studies modeled skull, brain, and CSF [12, 13] with coarse meshing to run their models with available computing power. However, they sacrificed important anatomical details of head structures (such as brain sulci and gyri structure or finer details of the cranium surface). Thereby, they failed to capture comprehensive details of the head responses. Recent advancements in computing technology enabled researchers to generate high-quality finer meshes to represent those complex anatomical details [9, 14, 15]. Nonetheless, a finer mesh can be time-consuming. Therefore, it is essential to define the optimized mesh size that provides time-effective simulations while, at the same time, do not compromise real-world anatomical fidelity. Another crucial aspect of meshing is the choice of the element type [16]. Although tetrahedral elements have showcased their efficacy in meshing complex geometries, they encounter volume locking issues when applied to nearly incompressible materials like the brain and CSF [17]. This challenge can be mitigated through the adoption of higher-order tetrahedral elements [16, 18]; However, higher order tetrahedral elements can increase the computational cost of the model. In contrast, the implementation of hexahedral elements presents an attractive option for meshing brain structures. Although capturing the intricate nuances of human brain geometry can be challenging with their use [19], hexahedral elements have consistently proven efficient in modeling brain structures. Importantly, they do not suffer from the volume locking issues associated with lower-order tetrahedral elements. This makes hexahedral elements an attractive option when balancing the need for accuracy and computational efficiency in brain meshing endeavors.

Despite these challenges, many computational FE models have been developed over the years to investigate the causation and effectuation of head and brain injuries in various impact scenarios. These models differ greatly in anatomical details, ranging from low to high fidelity [15, 20] and from geometrically and mechanically simplistic [21] to complex [15] descriptions of various head and neck structures. Early head FE models were developed in 1970s [21, 22] wherein researchers used linear elastic material properties and simplified geometries to represent complex head tissues so that they could be solved with available, limited computational power. As more material properties and computing power gradually became available, many researchers attempted to add complexities to their head models. For instances, head FE models like the Kungliga Tekniska Hogskolan Royal Institute of Technology (KTH) model [23], Strasbourg University FE Head Model (SUFEHM) [24], University College Dublin brain trauma model (UCDBTM) [25], and Worcester Head Injury Model (WHIM) [26] have evolved into more sophisticated models with finer meshing and geometries over the course of time. The latest enhancements of these head FE models [27–29] involve incorporating detailed brain tissue structures and complex properties, such as brain anisotropy. However, these models did not include a detailed neck structure.

As the neck serves as the primary structural connection between the head and the rest of the body and provides stability, mobility, and load-bearing support to the head, ignoring the neck would neglect its important role in distributing forces and absorbing impacts during traumatic head impacts [30–32]. Consequently, several studies attempted to include various neck structures, such as cervical vertebra [33], cervical discs [34], and neck muscles and ligaments [15] in their head models. However, modeling the intricate geometric and mechanical aspects of head-neck structures presents a formidable challenge, demanding significant time, labor, and advanced methodology. Only a select group of researchers developed a comprehensive and sophisticated head-neck FE model. Notably, pioneering efforts such as the Global Human Body Model Consortium [35] and the Total Human Model for Safety [12], were specifically created to delve into the biomechanics of head impacts in motor vehicle accidents and include a multitude of anatomical features pertaining to the head and neck. However, these models particularly implement a coarse meshing of the brain (average mesh size ∼ 3mm). Nonetheless, despite this limitation, they remain some of the most widely employed tools for gaining insights into the biomechanics of the head and neck across various head and brain injury investigations [31, 36-38]. Recently, Tse et al. [39] and Tuchtan et al. [34] proposed head-neck FE models without active neck muscles. whereas, two recent studies included active neck muscles in their head-neck FE models [15, 40]. However, Liang et al. [15] omitted CSF from their head model and Barker and Cronin [41] model did not include brain in their model.

Furthermore, the advancement in medical imaging techniques, such as computed tomography (CT) and magnetic resonance imaging (MRI), has contributed to the development of biofidelic head-neck models. CT imaging and MRI scans are traditionally used to capture hard (e.g., vertebrae, skull, etc.) and soft tissues (e.g., muscles, ligaments, etc.), respectively. Some previous studies have used both CT and MRI techniques to image the irregular shapes of head and neck structures [15]. Nonetheless, it could be cumbersome to avoid alignment issues between CT and MRI images during the 3-D model development stage. In addition, CT imaging exposes human subjects to harmful radiation [42], whereas MRI uses strong magnetic fields to capture the tissues of interest without any ionizing radiation. Though MRI scanning is traditionally implemented to image soft tissues, the recent development of MRI techniques [43] has unlocked the door for high-quality imaging of both soft and hard tissues. To our knowledge, three previous studies [9, 14, 44] have exclusively used MRI datasets to develop a head FE model; however, none of them considered any neck structures.

Therefore, this study aimed to develop an MRI-based detailed head-neck FE model and validate its’ anatomical accuracy and biomechanical performance by using widely used experimental data. In comparison to existing head-neck FE models in the literature, the proposed head-neck model was developed with several methodological innovations: 1) the detailed head (composed of scalp, skull, dura and pia maters, CSF, gray and white matters, etc.) and neck (consisted of cervical spine, disc, ligaments, and muscles) structures of the model was developed by using only MRI scan data from a male participant (52^nd^ percentile by stature and 92^nd^ percentile by weight) with larger body mass index; 2) advanced data analysis method was implemented to accurately align and interact unscaled adjacent model geometries; 3) a novel meshing method was applied to generate an unstructured, finer hexahedral structure of the brain’s gray and white matters and the CSF (average mesh size ∼ 1 mm); and 4) an erosion model was implemented to the scalp to simulate its realistic energy absorption capability under any given mechanical impact.

## Materials and Methods

### MRI-derived head-neck FE model development

The methodological framework of our head-neck FE model development procedures is presented in Figure 1. We used head and neck MRI datasets of a male firefighter (age: 42 years, height: 176 cm, weight: 106 kg, BMI: 34.2 kg/m^2^) throughout the whole modeling procedure. The head anthropometric measures and percentile distribution of the participant is presented in Table 1. Prior to the MRI procedure, we collected written informed consent from the subject. The study protocol was approved by the local institutional review board (*IRB 2020-708*).

**Fig. 1.**
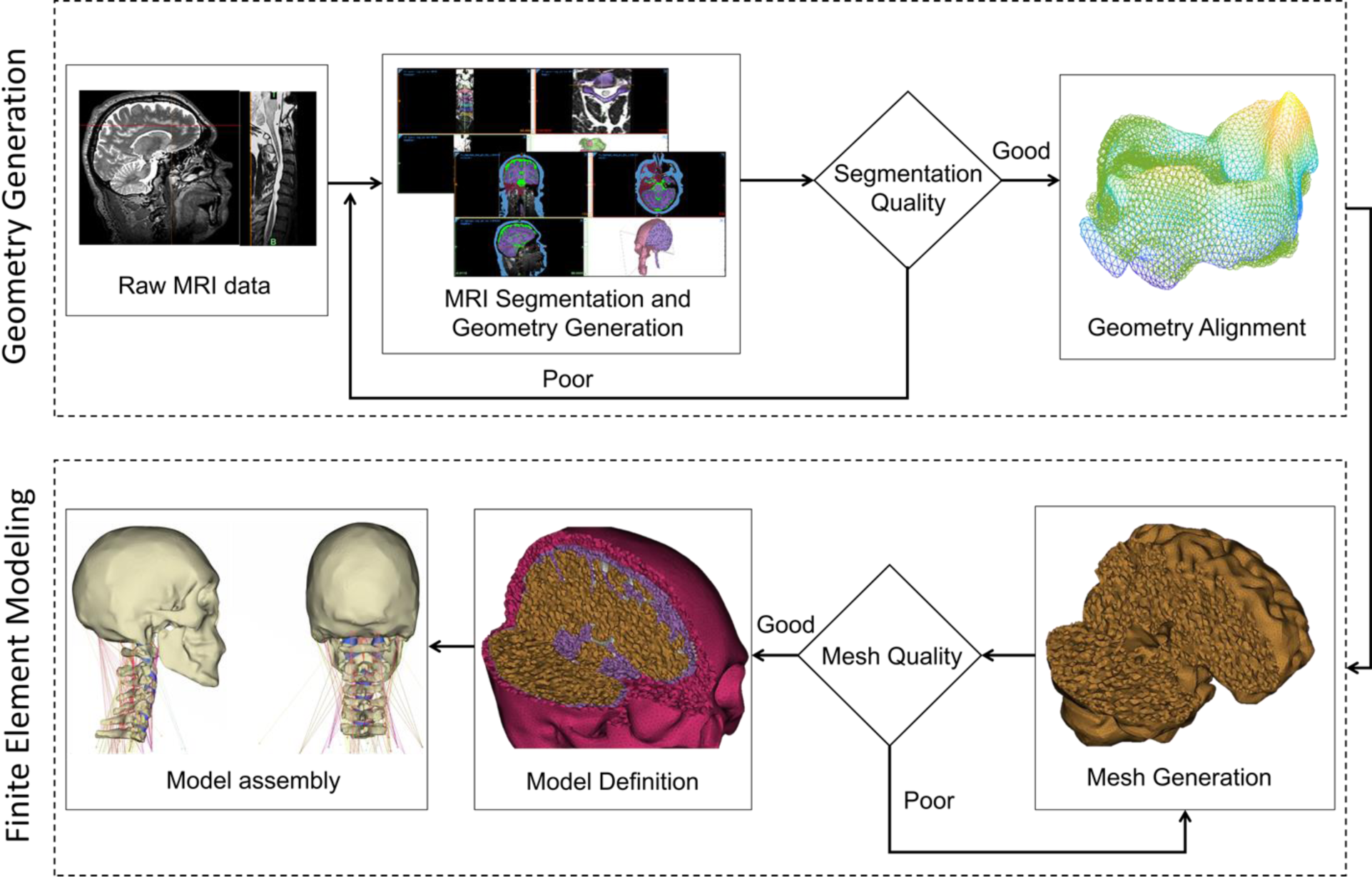
A schematic presentation of the methodological framework. The framework follows four consecutive steps: 1) 3-D head-neck geometry development from MRI datasets, 2) Geometrical verification of developed images, 3) FE meshing of the model geometries, and 4) defining material properties. Steps to generate and verify head-neck geometry are shown in the top dashed box, and the steps for the finite element modeling are provided in the bottom dashed box.

**Fig. 2.**
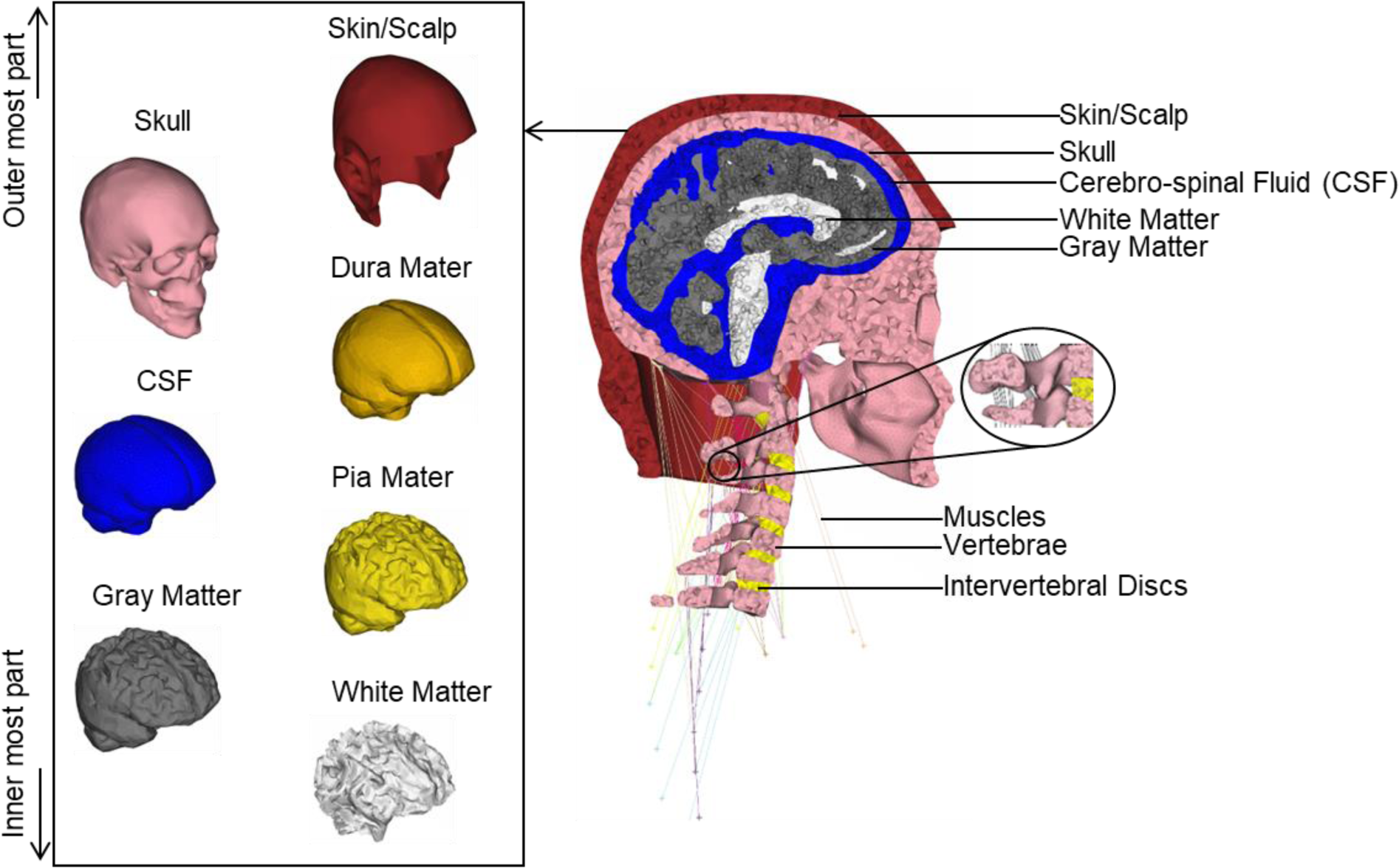
A schematic presentation of our developed head-neck finite element model consisting of scalp, skull, brain (gray and white matter), CSF, pia mater, dura mater, cervical vertebrae (C1-C7), intervertebral discs, muscles (obliquus capitis superior, superior longus colli, rectus capitus major, rectus capitus minor, longus capitis, rectus capitis ant, rectus capitis lat, anterior scalene, middle scalene, posterior scalene, sternocleidomastoid, longissimus capitis, longissimus cervicis, multifidus cervicis, semisplenius capitus, semispinalis cervicis, splenius capitis, splenius cervicis, levator scapula, oblique capitus inferior and trapezius), and ligaments (anterior longitudinal ligament, posterior longitudinal ligament, ligamentum flavum, capsular ligament, interspinous ligaments, tectorial membrane, anterior and posterior atlanto-occipital ligaments, anterior and posterior atlanto-axial ligaments, apical ligament, alars ligament, transverse ligament, and cruciate ligament of atlas)

**Table 1.** Anthropometric measures of the study participant and percentile distribution with regard to ANSUR II database [45].

|  | Value | ANSUR II percentile |
| --- | --- | --- |
| Stature (cm) | 176.0 | 52<sup>nd</sup> |
| Weight (kg) | 106.0 | 92<sup>nd</sup> |
| Head length (cm) | 20.0 | 54<sup>th</sup> |
| Head circumference (cm) | 57.8 | 62<sup>nd</sup> |
| Head breadth (cm) | 16.0 | 88<sup>th</sup> |

### Head-neck geometry development

We obtained hard and soft tissue images of head-neck structures ranging from the top of the scalp to the third thoracic vertebra by implementing T-1 weighted and T-2 weighted sequences at a 3T MRI scanner (Siemens Medical Solution, Germany). T2-weighted images were primarily used since this sequence employed different gray values to distinguish soft and hard tissues and fluids more accurately. Following image acquisition, we meticulously processed the MRI data using MIMICS 24.0 and 3-Matic 16.0 software platforms (both from Materialize Inc., Belgium). To enhance the contrast between soft and hard tissues in the neck region, we resliced the MRI data at 0.5 mm intervals. Subsequently, we employed identifiable masks to segment the MRI images of each head-neck structure of interest and converted them into three-dimensional (3-D) representations using MIMICS. For the segmentation of the brain, we utilized the MATLAB’s statistical parametric mapping (SPM) 12 toolbox to segment CSF and brain’s gray and white matter. To ensure the accuracy and fidelity of our 3-D geometries, we transferred the 3-D point cloud data of each structure into the 3-Matic platform, to remove additional unwanted noise and tissues that were not eliminated during segmentation in MIMICS (Fig. 1).

### Head-neck geometry verification

Once the 3-D geometries of all head-neck structures were created, we verified the accuracies of their individual geometrical parameters (e.g., shape, size, and volume) and alignments with the literature data [46–50]. These studies were selected due to the use of live healthy adult subjects in their works. Their geometrical parameters were determined by following steps as described in previous studies [46–50]. The scalp thickness (minimum) was automatically calculated in the ANSA (BETA CAE Systems SA, Greece) software, whereas the maximum scalp thickness was chosen as the highest thickness value among the values manually measured at frontal, temporal, occipital, and parietal lobes [48]. The average skull thickness was measured in six cranium sites (F3, F4, T3, T4, P3, P4) as specified by [48]. Brain and CSF volumes were also measured automatically in the ANSA. The geometries of neck vertebral bodies and discs were measured by identifying four corner-most points (anterior-superior, anterior-inferior, posterior-superior, posterior-inferior points) of the vertebral body and two corner-most points (distal and proximal ends) of their spinous process in the mid-sagittal plane as described in the literature [50]. The vertebral height was the average of anterior (distance from anterior-superior to anterior-inferior points) and posterior (distance from posterior-superior to posterior-inferior points) body heights, the vertebral depth was the average of superior (distance from posterior-superior to anterior-superior points) and inferior (distance from anterior-inferior to posterior-inferior points) vertebral body widths, spinous process length was the distance between distal and proximal corner-most points of the spinous process, and the vertebral body to spinous process length was the distance between the distal end of the spinous process and posterior side of the vertebrae body. Furthermore, we calculated angle-corrected disc heights as described in literature [51].

During the MRI procedure, the subject was required to lie flat and use pillows to immobilize his head and neck, which might have led to some degree of neck tilt. Thence, we examined the orientation of each neck vertebral body with respect to a neutral, upright neck orientation. We employed principal component analysis to calculate principal components and the centroid of each vertebral body in MATLAB R2021b (MathWorks, USA) platform. Then, we rotated each vertebral body with regard to its own centroid and matched its’ first three principal components with the global coordinates of the model. Furthermore, we rotated and translated C2, C3, C4, C5, C6, and C7 vertebrae to match the frontal and sagittal planes of the C1 vertebra.

### Mesh generation and model assembly

We used the ANSA platform to generate highly detailed, fine meshes for all structures within the head-neck region, avoiding any scaling. Upholding the mesh quality was of paramount importance to us. Therefore, we set stringent threshold values of 4 for aspect ratio, 45 for skewness, 20 for warping, and 140 for the maximum angle during the mesh generation process. The resulting meshed model comprised of 1.35 million elements and the mesh structures of individual structures are provided in Table 2. For the regions encompassing nearly incompressible tissues such as grey matter, white matter, and CSF, we adopted a two-step meshing approach. Initially, a tetrahedral meshing algorithm was employed to create an unstructured tetrahedral mesh, accurately representing the anatomical features of these regions. Subsequently, a custom script was utilized to convert each tetrahedral element into four hexahedral elements, resulting in a purely unstructured hexahedral mesh for these areas. Briefly, our custom script employed an algorithm to connect midpoints of each tetrahedral element edge to the midpoint of each tetrahedral element face, splitting each tetrahedral element into four hexahedral elements. To address potential element distortion, we applied stricter quality criteria (2 for aspect ratio, 30 for skewness, 10 for warping, and 120 for the maximum angle) for the original tetrahedral elements. Nevertheless, hexahedral elements that did not meet this stringent quality criteria were manually fixed. This novel methodological approach for hexahedral meshing of the brain and CSF tissues was very user-friendly (automatic and less time-consuming), and thus can facilitate researchers to address one of the challenges during detailed brain-CSF FE model development process.

**Table 2.** The mesh structure details of individual head-neck components.

| Model part | Number of elements | Element type | Aspect ratio < 3 |
| --- | --- | --- | --- |
| Scalp | 267720 | 3-D Tetrahedral | 100% |
| Skull | 230473 | 3-D Tetrahedral | 99% |
| Neck | 71894 | 3-D Tetrahedral | 99% |
| Dura mater | 36804 | 2-D Quad shell | 80% |
| Pia mater | 13536 | 2-D Quad shell | 84% |
| CSF | 138316 | 3-D Hexahedral | 88% |
| Grey matter | 360240 | 3-D Hexahedral | 85% |
| White matter | 236188 | 3-D Hexahedral | 94% |
| Muscles | 132 | 1-D beam | - |
| Ligaments | 486 | 1-D beam | - |

Furthermore, we included 42 muscles on both left and right sides of the neck (21 on each side) and 14 ligaments around the neck. We modeled the ligaments as linear springs with 486 1-D beam elements, and their stiffness was adopted from literature [52]. Similarly, neck muscles were also modeled as 1-D beam elements, with a total of 132 elements. The origin and insertion points of these muscles and ligaments were adopted from Gray’s anatomy book [53].

### Defining material properties and model assembly

The material properties and governing equations of all head-neck structures included in our model are provided in Table 3. Briefly, we modeled skull, pia mater, dura mater, and vertebrae as linear elastic material [33, 34, 54], scalp as a linear viscoelastic material model [55], CSF as nearly incompressible one-term hyper-elastic material [56], intervertebral discs as a hyper-viscoelastic material model [57], and brain (gray and white matter) as a hyper-viscoelastic material [58, 59]. Previous literature [8] has shown that white matter is approximately 40% stiffer than grey matter, therefore we modeled the grey and white matters accordingly. Some previous head FE studies have used a Poisson’s ratio of 0.49999 in their brain model [16, 18], which is close to the theoretical limit for near-incompressible materials. However, we used a Poisson ratio of 0.4977, derived from experimental material characterization data found in the existing literature specific to the brain [59], and thus provided the brain a certain extent of compressibility property. As the scalp shows load-rate-dependent mechanical characteristic [5] and has very low stiffness, it is expected to rupture under high-impact loads. Therefore, we applied the erosion model available in the LS-DYNA (Livermore Software Technology Corporation, USA) platform to identify and remove ruptured tissues if the strain of any scalp element exceeded 60% [60]. Moreover, we modeled the muscles using the Hill-type muscle model. Muscle volume and physiological cross-section area (PCSA) were taken from the literature [61]. Muscle resting length was calculated as muscle volume divided by its’ PCSA. The maximum velocity of a muscle was set to 10 times its resting length. The maximum stress of a muscle was set to 0.3 MPa [62].

**Table 3.** Material properties and constitutive equations of each head-neck structure included in our model.

| Segment | Material model | Material constants | Constitutive Equation |
| --- | --- | --- | --- |
| Scalp [55] | Linear viscoelastic | $G_0 = 1.70 \text{ MPa}$<br>$G_\infty = 0.68 \text{ MPa}$<br>$K = 20 \text{ MPa}$<br>$\beta = 0.00003$<br>$\rho = 1100 \text{ kg/m}^3$ | $G(t) = G_\infty + (G_0 - G_\infty)e^{-\beta t}$ |
| Skull[54] | Linear elastic | $E = 15.00 \text{ GPa}$<br>$\rho = 1800 \text{ kg/m}^3$<br>$\nu = 0.21$ | $\sigma = E\varepsilon$ |
| Dura mater [34] | Linear elastic | $E = 5.00 \text{ MPa}$<br>$\rho = 1200 \text{ kg/m}^3$<br>$\nu = 0.45$ | $\sigma = E\varepsilon$ |
| Pia mater [34] | Linear elastic | $E = 2.30 \text{ GPa}$<br>$\rho = 1000 \text{ kg/m}^3$<br>$\nu = 0.45$ | $\sigma = E\varepsilon$ |
| Vertebrae [33] | Linear elastic | $E = 8.00 \text{ GPa}$<br>$\rho = 1200 \text{ kg/m}^3$<br>$\nu = 0.22$ | $\sigma = E\varepsilon$ |
| CSF[56] | Hyper-elastic | $C_{10} = 0.0112$<br>$\rho = 1000 \text{ kg/m}^3$<br>$\nu = 0.499$ | $W(J_1, J_2, J) = \sum_{p,q=0}^n C_{pq}(J_1 - 3)^p (J_2 - 3)^q + W_H(J)$ |
| White matter [58, 59] | Hyper-viscoelastic | $\mu_0 = 1.18 \text{ kPa}, \alpha = -4.77$<br>$\rho = 1060 \text{ kg/m}^3$<br>$\nu = 0.4977$<br>$g_1 = 0.653$<br>$g_2 = 0.206 \text{ M}$<br>$\tau_1 = 0.448 \text{ s}, \tau_2 = 15.007 \text{ s}$ | $W = \frac{2\mu_0}{\alpha^2} (\bar{\lambda}_1^\alpha + \bar{\lambda}_2^\alpha + \bar{\lambda}_3^\alpha - 3)$<br>$\mu = \mu_0 \left[ 1 - \sum_{i=1}^2 g_i (1 - e^{-t/\tau_i}) \right]$ |
| Grey matter [58, 59] | Hyper-viscoelastic | $\mu_0 = 0.84 \text{ kPa}, \alpha = -4.77$<br>$\rho = 1060 \text{ kg/m}^3$<br>$\nu = 0.4977$<br>$g_1 = 0.653$<br>$g_2 = 0.206 \text{ M}$<br>$\tau_1 = 0.448 \text{ s}, \tau_2 = 15.007 \text{ s}$ | $W = \frac{2\mu_0}{\alpha^2} (\bar{\lambda}_1^\alpha + \bar{\lambda}_2^\alpha + \bar{\lambda}_3^\alpha - 3)$<br>$\mu = \mu_0 \left[ 1 - \sum_{i=1}^2 G_i (1 - e^{-t/\tau_i}) \right]$ |
| Intervertebral discs [57] | Hyper-viscoelastic | $C_{10} = 0.15, C_{01} = 0.03$<br>$\rho = 1000 \text{ kg/m}^3$<br>$\nu = 0.499$<br>$G_1 = 1.70, G_2 = 1.20, G_3 = 2.00$<br>$\tau_1 = 11.76, \tau_2 = 1.10, \tau_3 = 0.13$ | $W(J_1, J_2, J) = \sum_{p,q=0}^n C_{pq}(J_1 - 3)^p (J_2 - 3)^q + W_H(J)$<br>$G(t) = \sum_{k=1}^n G_k \exp^{-t/\tau_k}$ |
| Muscles [61, 62] | Hill-type | PCSA[61]<br>Muscle volume[61] | $F_{active} = F_{max} \times f_{FL} \times f_{FV} \times A(t)$<br>$F_{Passive} = \frac{F_{max}}{e^{K_{sh}} - 1} \left[ e^{\frac{K_{sh}}{L_{max}} \left( \frac{L}{L_{rest}} - 1 \right)} - 1 \right] \text{ For } L > L_{rest}$ |
| Ligaments [52] | Linear spring | k[52] | $F = kx$ |

E: Elastic modulus, $G_0$ : short-term shear modulus, $G_\infty$ : long-term shear modulus, K: bulk modulus, $\beta$ : decay constant, $\rho$ : density, $\nu$ : Poisson ratio, $\tau_k$ : relaxation time, $g_k$ : relaxation coefficient, $\bar{\lambda}_i^\alpha$ : principal stretch, W: strain energy density function, $C_{pq}$ , $\mu$ : shear modulus, $\alpha$ : material constants, $F_{active}$ : active muscle force, $F_{passive}$ : passive muscle force, $f_{FL}$ : force-length relation of the muscle, $f_{FV}$ : force-velocity relation of the muscle, $A(t)$ : muscle's activation level versus time relationship, $L_{rest}$ : the resting length and $K_{sh}$ : passive muscle force constant, k: ligament stiffness.

Similar to previous studies [9, 55], we implemented tied boundary contact between discs and vertebrae, skull and scalp, Skull and dura mater, dura mater and CSF, CSF and pia mater, brain and pia mater and neck-skull articulations. Moreover, the bottom surface of the C7 cervical vertebra was fixed to restrict its’ all degrees of freedom.

### Experimental Data for head-neck FE model validation

Four head impact experiments [63–66] were used to validate the efficacy of our developed head-neck FE model (Fig. 3). The first experimental dataset was the linear acceleration profile (Fig. 3a) of an in-vivo study conducted by the Naval biodynamics laboratory (NBDL) [67] in 1972. [63] corrected NBDL’s linear acceleration profile for T1 vertebral rotation in 1995. We applied one of those corrected acceleration profiles [63] (Fig. 3c) to our whole head-neck model. As the subject’s neck muscles were pre-tensed in the NBDL experiment, we modeled the flexor muscles with 10% activation and the extensor muscles with 80% activation.

**Fig. 3.**
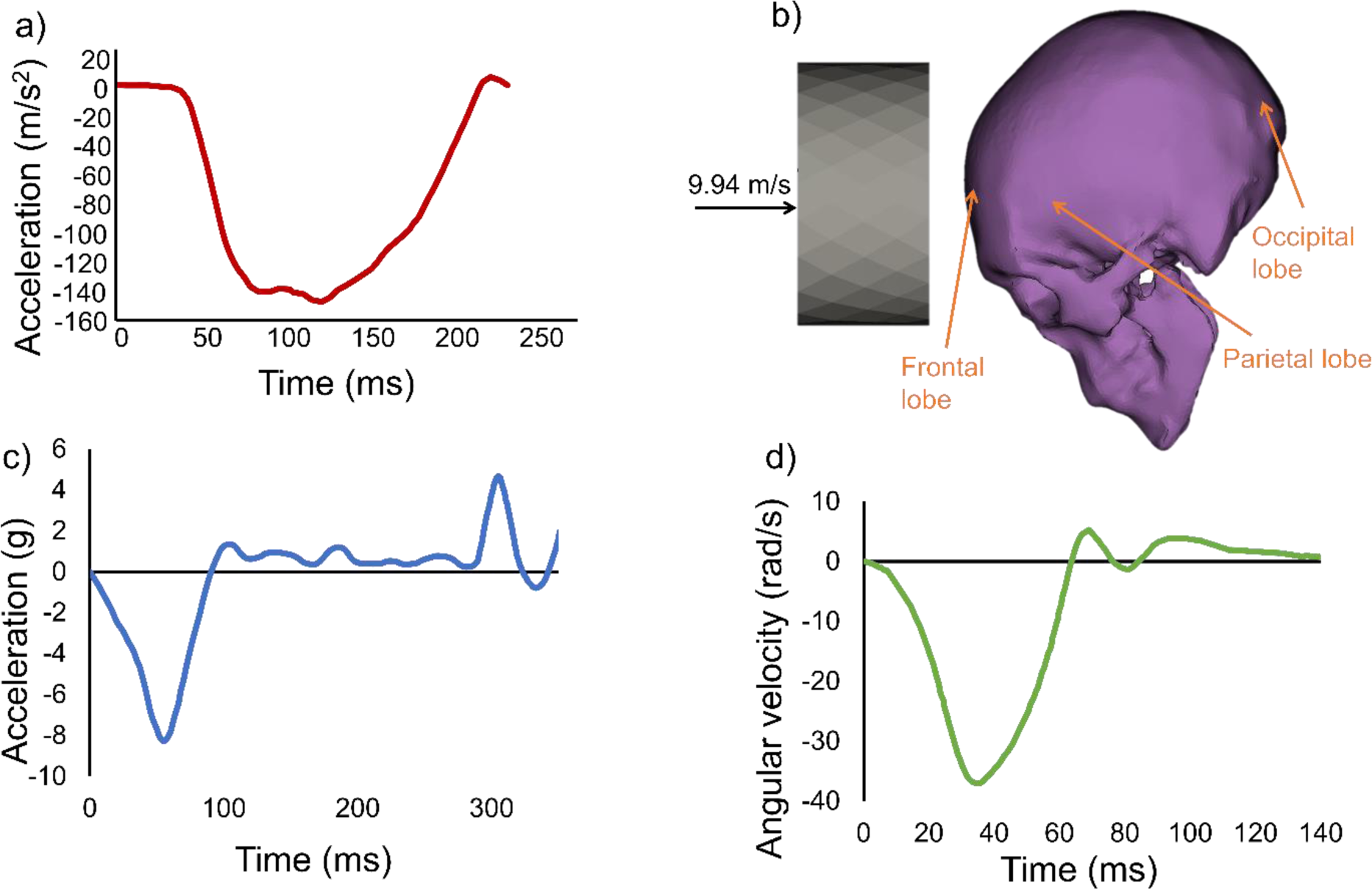
Experimental impact scenarios that were numerically replicated in this study: a) linear acceleration profile of NBDL study [67] with respect to T1 vertebral rotation [63], and b) Nahum et al.[64], in which they hit a cadaver head using rigid body impactor [64]. c) linear accleration profile for Ito’s [65] cervical vertabrae study, and d) Angular velocity profile for Alshareef ‘s [66] sonomicrometry study. The run time for finite element simulations of NBDL,’ Nahum’s, Ito’s, and Alshareef’s studies were respectively 220 ms, 10 ms, 350 ms, and 150 ms.

Our second experimental dataset was the impact study conducted by [64], in which they analyzed the mechanical response (intracranial pressure) of the head of a cadaver subject. Similar to their experimental condition, we simulated a rigid aluminum impactor with a mass of 5.59 kg and a velocity of 9.94 m/s blowing the frontal bone of the skull in the direction of mid-sagittal plane, while the head was tilted 45° downward towards the ground (Fig. 3b). We meshed the impactor with 174,596 tetrahedral elements. The impact pulse lasted for about 5 ms, with a peak impact force of 9.41 kN at around 2.5 ms. We replicated Nahum’s study for two model conditions. As the Nahum’s study was originally performed using head-only cadaver, we first implemented the head-only model to validate the simulation results of our developed model. Then, we replicated the Nahum’s study condition for the whole head-nek model to explore the differences in numerical outcomes when considering neck structures.

Our third scenario involved data from Ito et al. [65] cervical vertebrae study to validate how our modeled neck responds to linear impact in comparison to those with the experimental neck cadaver. To do so, we applied their linear acceleration profile to the C7 vertebrae (Fig. 3c), where they subjected the neck to an 8g horizontal acceleration and measured disc strain. To replicate their muscle force simulation, which employed 4.0 N/mm springs for anterior and lateral springs and 8.0 N/mm for posterior springs, we adjusted our model with an 80% activation level for neck extensor muscles and a 40% activation level for neck flexor muscles.

Lastly, our fourth experiment drew from Alshareef et al. [66] sonomicrometry study. They measured brain displacement in response to a 40 rad/s, 60 ms, angular velocity (Fig. 3d) and reported three of their sensor data (receivers 9, 16, and 3) located in the parietal region of the brain. Although their specimen included the neck, the muscles were not active during the experiment. We represented the muscular condition with passive muscle representation (0% activation). We used LS-DYNA explicit solver platform installed in the Texas Tech university high-performance computing center (two AMD EPYC 7702 CPUs and 500 GB memory) to solve all impact simulations. A simulation of 220 ms (NBDL study) took about 36 hours to complete.

### Post-processing and statistical analysis

We used META (BETA CAE Systems SA, Greece) software for post-processing and retrieving numerical solutions of all four impact simulations. We assessed intracranial pressure (ICP) in frontal, parietal, and occipital lobes for both head-only and head-neck model simulations and compared our results against Nahum’s impact results. Additionally, we compared the von-Mises stress of the brain at three impact instances to explore the contribution of neck structures on mechanical responses of the brain to Nahum’s rigid body impacts. We also compared model-predicted neck flexion angle data and their peak time with the experimental neck flexion data of the NBDL study [63]. To compare with the actual experimental results of Ito’s study [65], we calculated cervical disc strain at every disc level. In addition, we conducted an evaluation of brain deformation at specific points highlighted in Alshareef’s study [66] to validate our own brain deformation findings. Pearson correlation analysis was employed for pairwise comparison between all numerical and experimental data patterns. For this purpose, we digitized Alshareef’s experimental brain deformation results [66], Nahum’s experimental ICP plots [64] and NBDL’s experimental neck flexion angle plot [63] in the MATLAB platform. For the same digitized points, we retrieved model-predicted ICP data and neck positional information in the META platform. The neck flexion angle was the angle between the vertical line and the line joining the anterior-most point of the foramen magnum and the anterior-inferior corner-most point of C7 in the mid-sagittal plane. Furthermore, it’s important to note that each of these experimental studies utilized distinct coordinate systems. To ensure a precise and meaningful comparison between our numerical findings and their experimental data, we transformed our results to align with the coordinate systems employed in each respective study.

## Results

### 3-D head-neck geometry

As our head-neck FE model was developed from MRI data, the movement of the subject—particularly breathing—during MRI scans, we made few minor adjustments to individual geometries during image processing stage. Thus, to confirm the geometric accuracy and structural integrity, we compared our model’s head-neck geometry data against experimentally measured values reported in the literature (Tables 4-6). The head geometry data exhibited that both skull thickness and brain volume of our model were within one standard deviation (SD) of the values reported in previous literature [47, 48] (Table 4). On the other hand, the model’s CSF volume was found to be larger than the literature data [46] – about 29.52% greater than the maximum reported value. The maximum scalp thickness value was also observed to be 25.70% higher than the literature data [49].

**Table 4.**
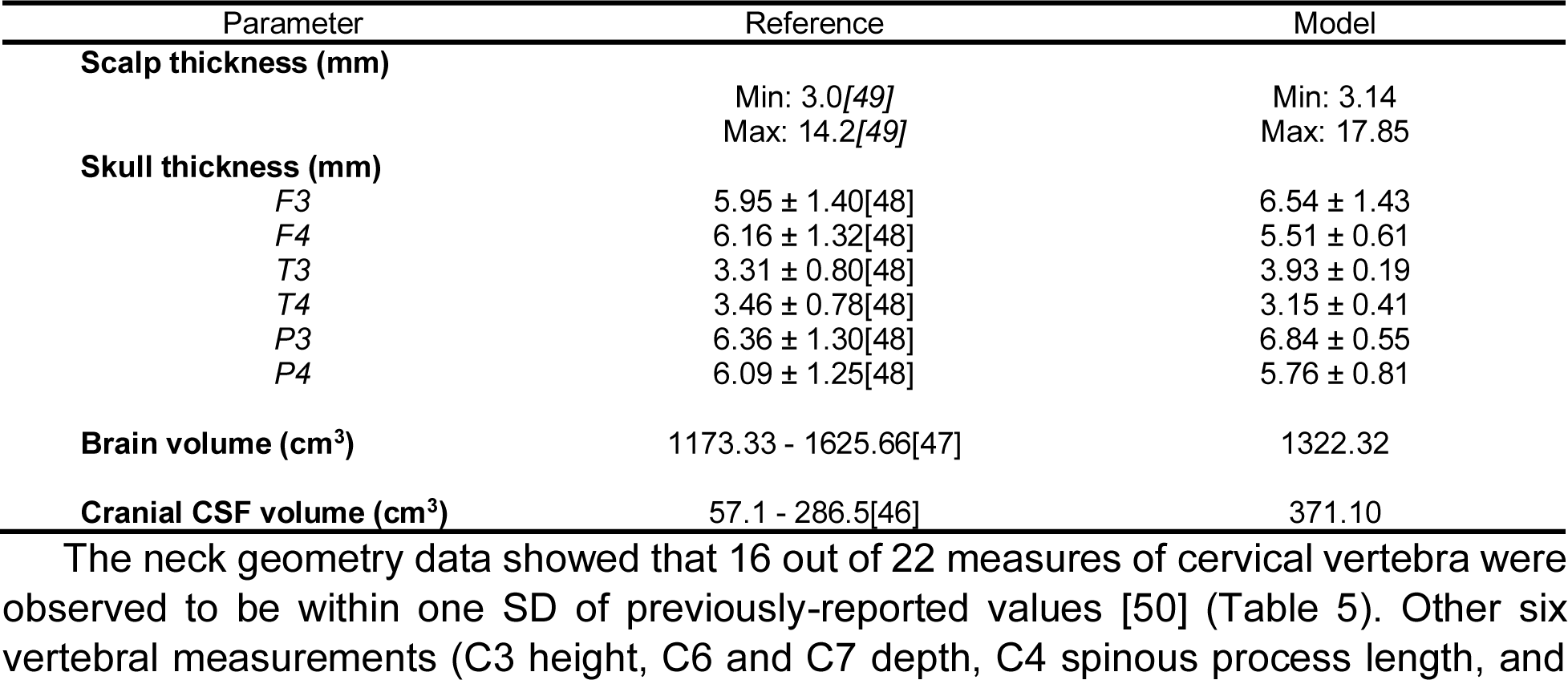
Comparison between the geometrical parameters of head structures and previously-reported experimental values. F3, F4, T3, T4, P3, P4 indicate skull measurement locations at frontal left, frontal right, temporal left, temporal right, parietal left, and right parietal skulls, respectively.

The neck geometry data showed that 16 out of 22 measures of cervical vertebra were observed to be within one SD of previously-reported values [50] (Table 5). Other six vertebral measurements (C3 height, C6 and C7 depth, C4 spinous process length, and C3 and C4 vertebral body to spinous process length) were within 17% of their corresponding reported values [50]. The disc height data also revealed that, except C5/C6 disc, all other discs were within one SD of the reported values [51] (Table 6).

**Table 5.**
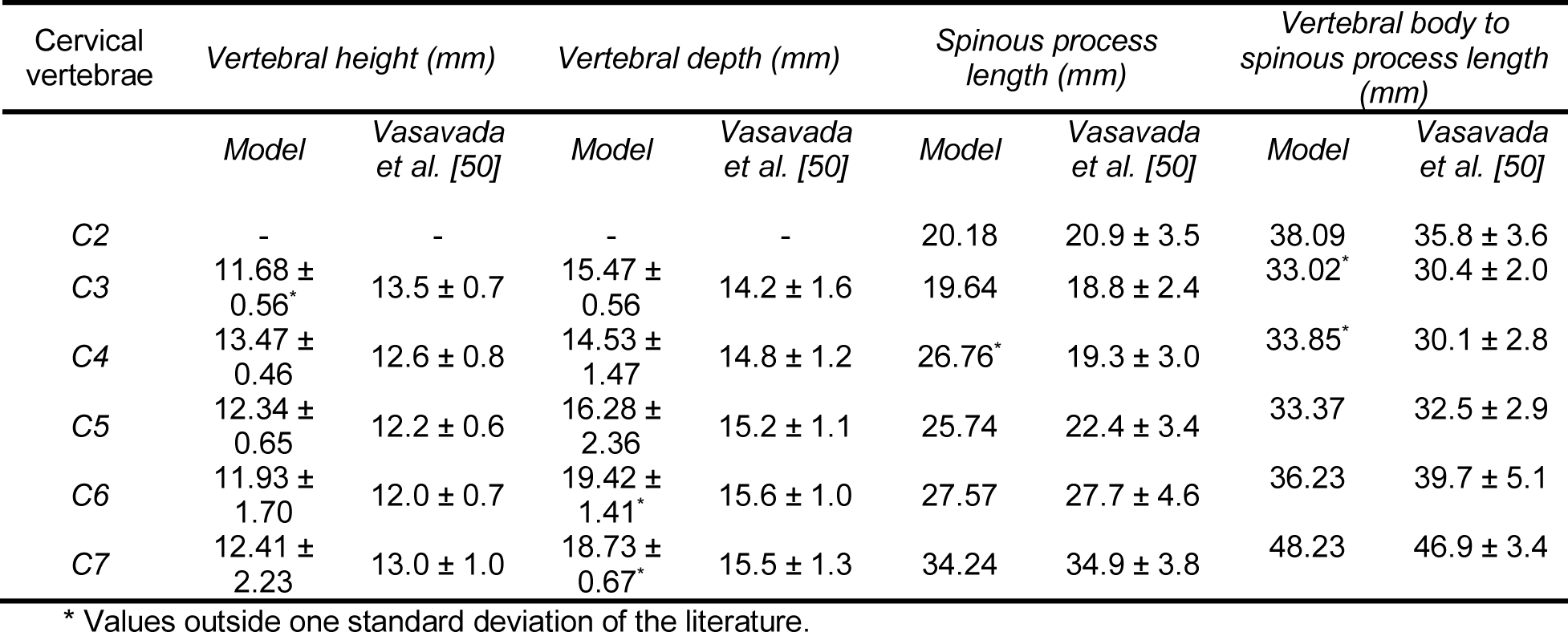
Comparison between cervical vertebral measurements and experimentally-measured values from literature.

| Cervical vertebrae | Vertebral height (mm) |  | Vertebral depth (mm) |  | Spinous process length (mm) |  | Vertebral body to spinous process length (mm) |  |
| --- | --- | --- | --- | --- | --- | --- | --- | --- |
|  | Model | Vasavada et al. [50] | Model | Vasavada et al. [50] | Model | Vasavada et al. [50] | Model | Vasavada et al. [50] |
| C2 | - | - | - | - | 20.18 | 20.9 ± 3.5 | 38.09 | 35.8 ± 3.6 |
| C3 | 11.68 ± 0.56* | 13.5 ± 0.7 | 15.47 ± 0.56 | 14.2 ± 1.6 | 19.64 | 18.8 ± 2.4 | 33.02* | 30.4 ± 2.0 |
| C4 | 13.47 ± 0.46 | 12.6 ± 0.8 | 14.53 ± 1.47 | 14.8 ± 1.2 | 26.76* | 19.3 ± 3.0 | 33.85* | 30.1 ± 2.8 |
| C5 | 12.34 ± 0.65 | 12.2 ± 0.6 | 16.28 ± 2.36 | 15.2 ± 1.1 | 25.74 | 22.4 ± 3.4 | 33.37 | 32.5 ± 2.9 |
| C6 | 11.93 ± 1.70 | 12.0 ± 0.7 | 19.42 ± 1.41* | 15.6 ± 1.0 | 27.57 | 27.7 ± 4.6 | 36.23 | 39.7 ± 5.1 |
| C7 | 12.41 ± 2.23 | 13.0 ± 1.0 | 18.73 ± 0.67* | 15.5 ± 1.3 | 34.24 | 34.9 ± 3.8 | 48.23 | 46.9 ± 3.4 |

\* Values outside one standard deviation of the literature.

**Table 6.** Comparison between angle-corrected, dimensionless intervertebral disc height of the model and their experimental counterparts.

| Cervical vertebral disc | Frobin et al.[51] | Model |
| --- | --- | --- |
| C2/C3 | 0.35 ± 0.07 | 0.303 |
| C3/C4 | 0.38 ± 0.08 | 0.366 |
| C4/C5 | 0.39 ± 0.06 | 0.375 |
| C5/C6 | 0.38 ± 0.04 | 0.295 |
| C6/C7 | 0.36 ± 0.06 | 0.323 |

## FE simulation results

### NBDL study

The kinematic responses of our head-neck model duplicated the NBDL’s experimental data very accurately. Our model showed a neck flexion angle of 36.45°±34.14°, with a peak of 88.31°, whereas the NBDL study reported 11 subjects’ neck flexion angles, ranged between a mean angle of 39.69°∼30.72° and the maximum angle of 67.20°∼87.50° [63]. Though a small number of discrepancies were found for the maximum neck angle (0.92% larger) and its’ peak time (5 ms delay), the visual comparisons of kinematic behavior at three-time instances revealed analogous head-neck responses between model predictions and experimental simulations (Fig. 4). In addition, Pearson’s correlation analysis evinced a strong positive correlation (r > 0.97) between experimental and numerical head-neck kinematic patterns (Fig. 4).

**Fig. 4.**
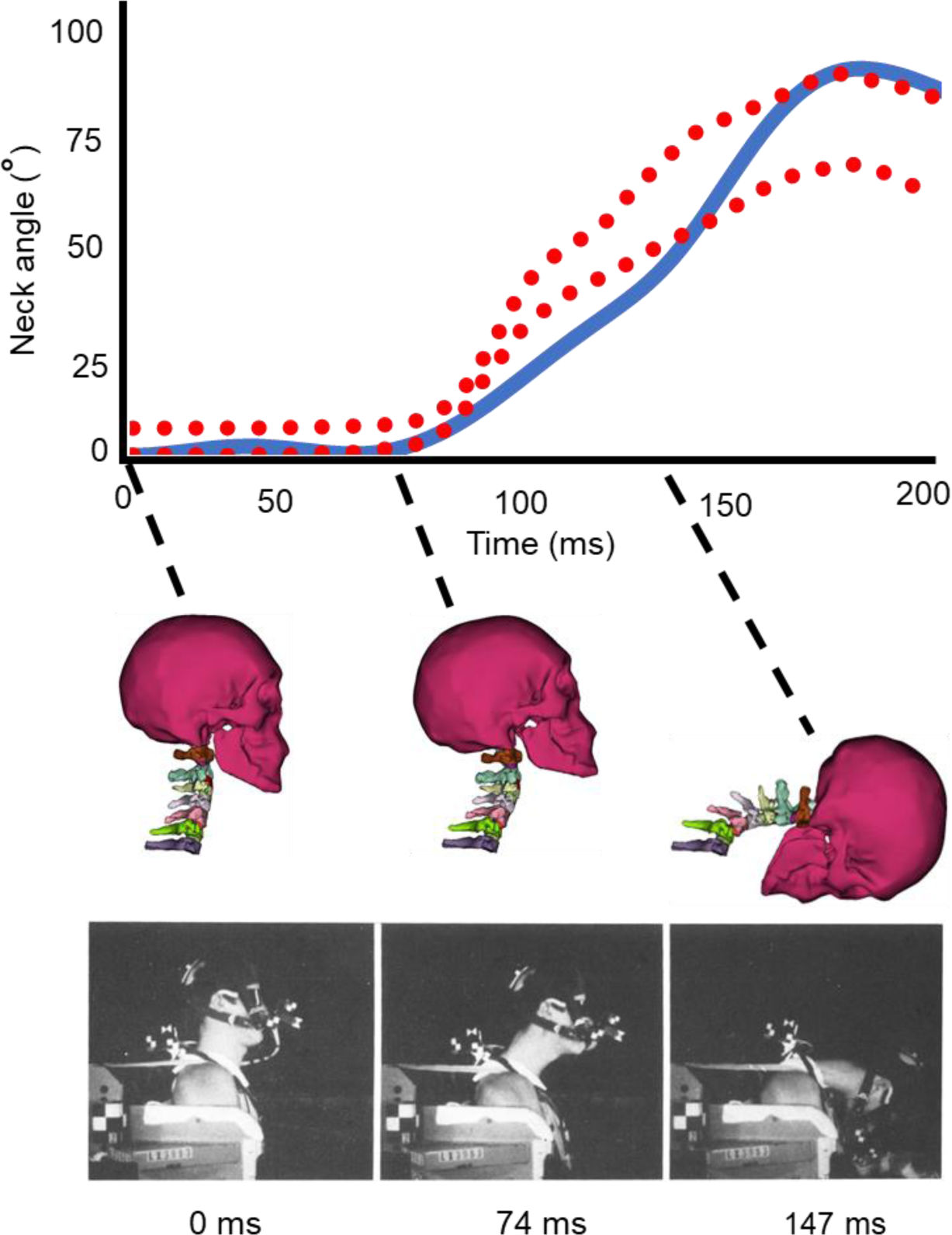
Comparison between model-predicted and NBDL-experimental neck flexion angles for the complete simulation duration (top) and a visual comparison between model-prediction (middle) and experimental head-neck kinematic responses (bottom) at three different time instances [63].

### Nahum study

We also found a good agreement (r = 0.80∼0.98) between model-predicted ICP data at frontal, parietal, and occipital regions for our head-only model and Nahum’s experimental (cadaver head) results, except for minor differences (less than 9%) in peak ICP values (Fig. 5a). Nonetheless, the maximum ICP differences in the frontal, parietal, and occipital regions for the head-only model, when compared to their corresponding experimental values, were 3.58%, 2.00%, and 8.14%, respectively. Conversely, the inclusion of neck structures resulted in significantly larger deviations, with ICPs differing by 13.47%, 16.61%, and 26.84% for the frontal, parietal, and occipital regions in comparison to the experimental data (Fig. 5b). Furthermore, a Pearson correlation analysis between the head-neck model predictions and Nahum’s experimental results revealed a relatively lower level of correlation (r = 0.66 ∼ 0.86) when compared to the head-only model results (Fig. 5a). The visual depiction of the impact scenarios in Figure 6 at three distinct time points illustrates that, across all three instances, the maximum von-Mises stress values within the brain were consistently lower in the head-neck model as compared to the head-only model.

**Fig. 5.**
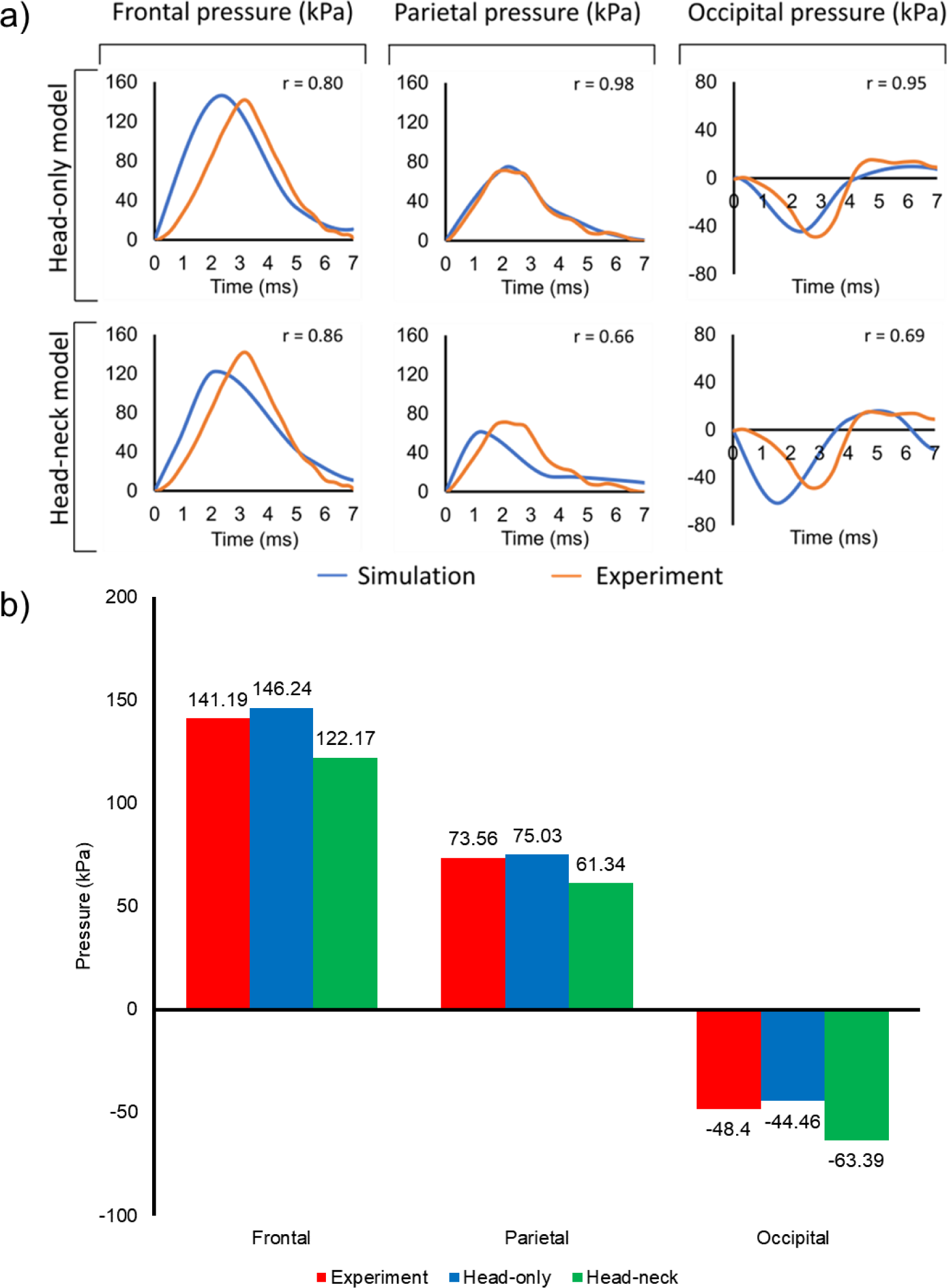
a) Comparison between simulated ICP results and Nahum’s experimental results [64] for both head-only (top) and head-neck (bottom) models at three different brain locations (frontal, parietal, and occipital). The correlation coefficients (r) are provided for each case. b) Maximum Absolute ICP value comparison between head-only and head-neck models and their experimental counterparts [64].

**Fig. 6.**
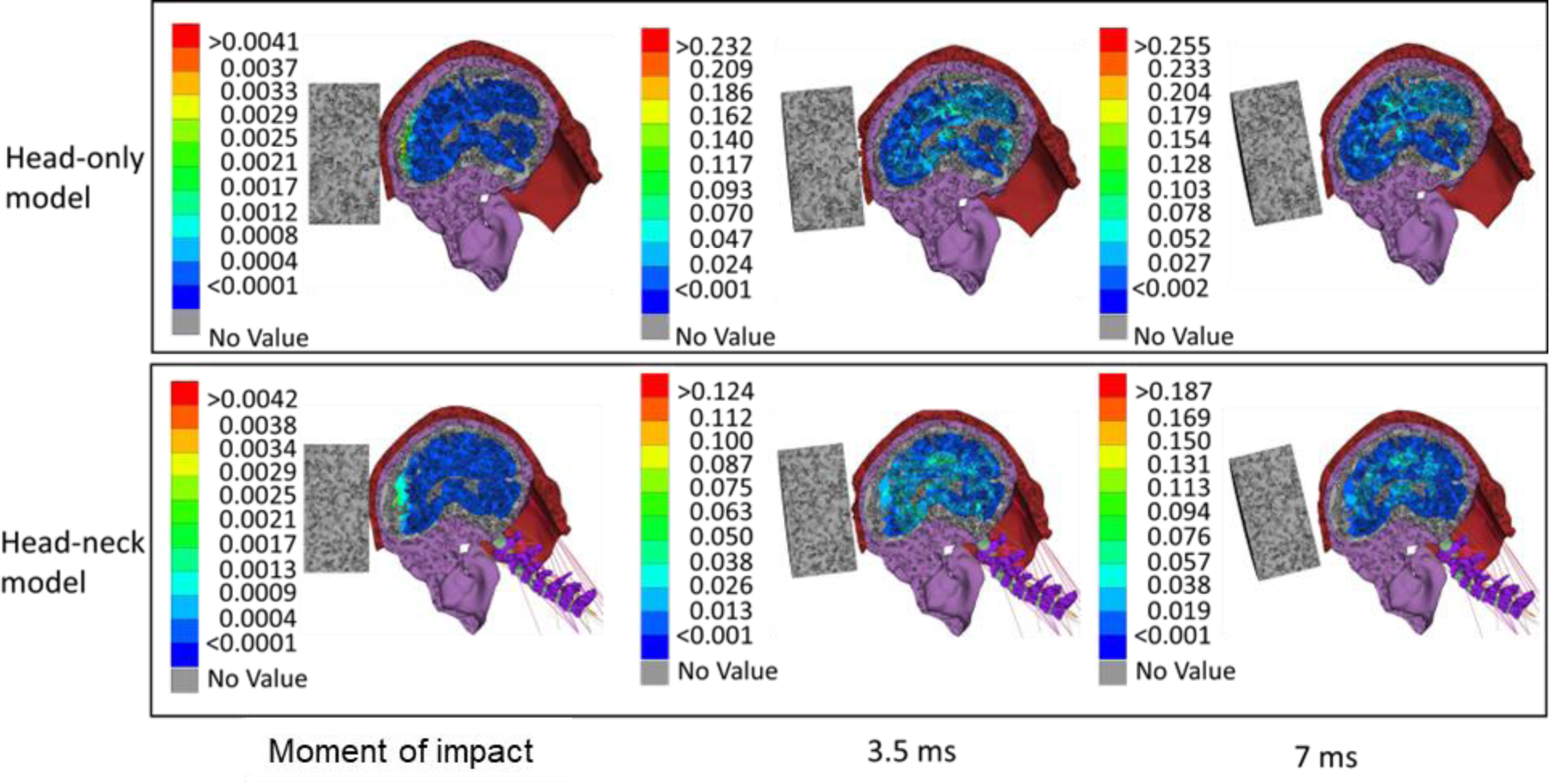
The overall von-Mises stress (MPa) of the brain. The color map and corresponding values indicate the range of von-Mises stress across all brain regions.

### Ito’s study

Our in-silico replication of Ito’s experimental study conditions [65] yielded peak shear strain values that were found to lie within one standard deviation of Ito’s experimental shear strain values for both anterior and posterior regions of individual intravertebral discs except the posterior region of the C2-C3 disc (Fig. 7) wherein the peak shear strain was more than one standard deviation but within the two standard deviations from its experimental counterpart. A notable difference in the peak shear strain values between experimental and simulation was also found in C2-C3 disc.

**Fig. 7.**
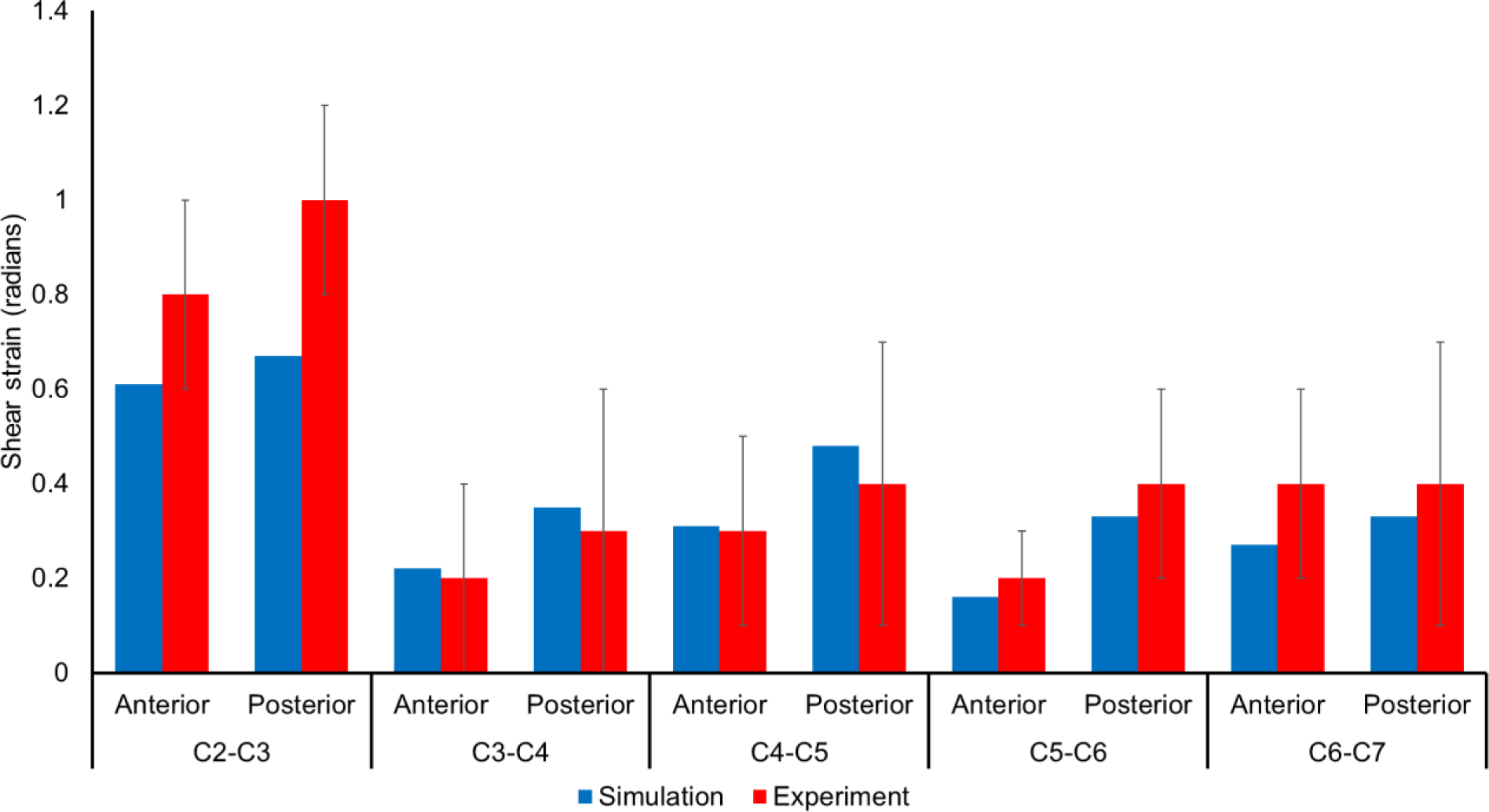
Comparison between simulated and experimental (mean ± SD) peak shear strain [65] in an 8g frontal impact scenario.

### Alshareef’s study

The brain displacement magnitudes of the parietal region of our model were compared with Alshareef’s three experimental sensor data placed in the parietal region (Fig. 8). The brain displacement patterns in the Y (medio-lateral) and Z (superior-inferior) directions at all three locations demonstrated strong agreement (r = 0.65 ∼ 0.91) with the experimental findings [66], as depicted in Figure 8a. In contrast, our displacement results in the X (anterior-posterior) direction displayed relatively a medium-level of correlation (r = 0.31 ∼ 0.58). As the displacements fluctuate around zero, computing a simple average of the brain displacement values would yield a value close to zero. Therefore, we compared the average of the absolute brain displacement and observed that slight disparities exist between numerical results and experimental data for receiver 9 in X (0.60%) and Z (7.46%) directions, receiver 16 in Y (2.62%) and Z (1.61%) directions, and receiver 31 in Y (8.18%) and Z (6.45%) directions. In contrast, A notable difference was found in the Y (45.75%), X (34.66%), and X (33.46%) directions for receiver 9, receiver 16, and receiver 31, respectively.

**Fig. 8.**
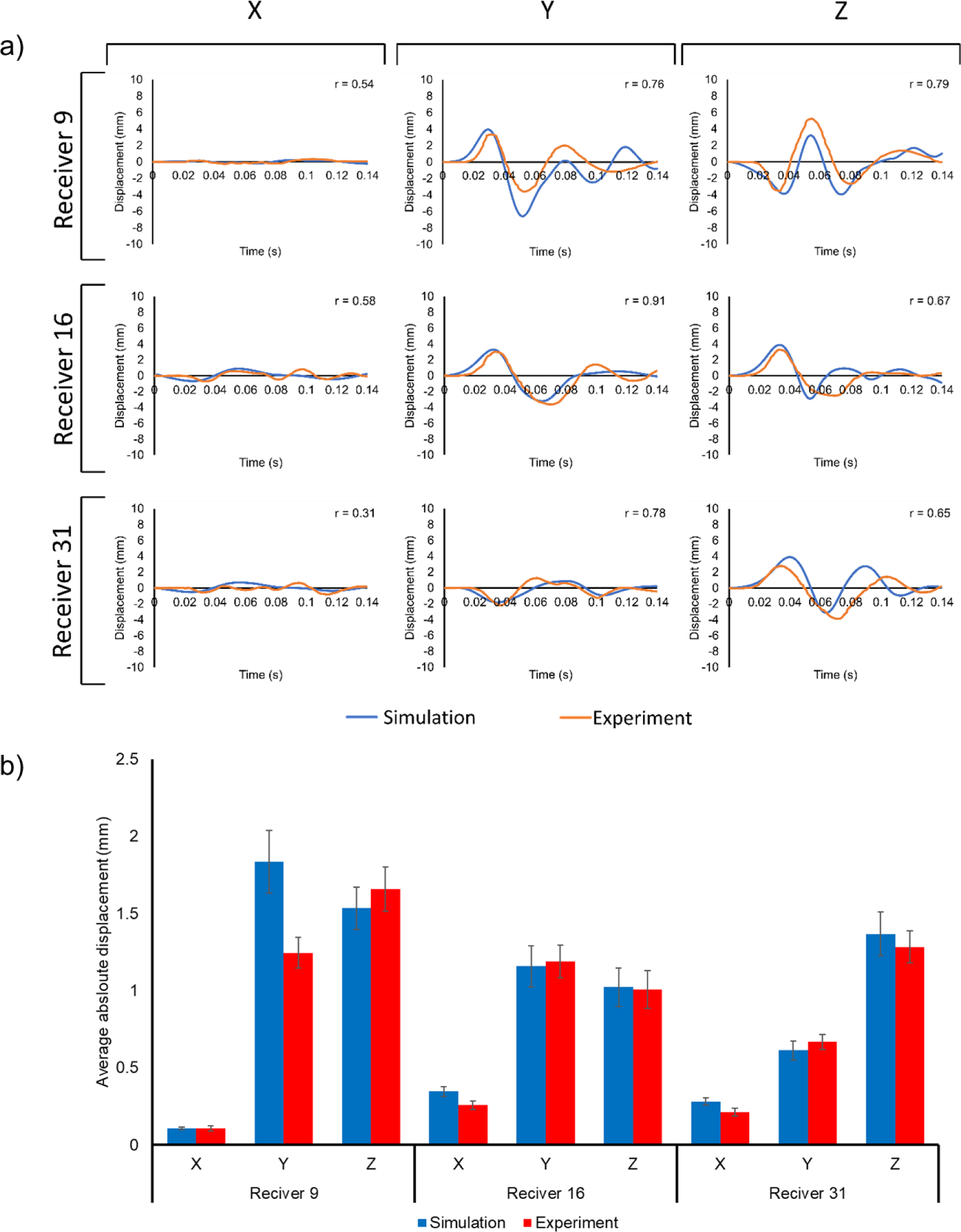
Comparative analysis of brain displacement between simulated and experimental results in X, Y, and Z directions across three receiver locations, as reported in the Alshareef et al. [66] study: a) the experimental and simulated brain displacement patterns and their corresponding correlation coefficients (r) in all three directions and b) the simulated and experimental absolute brain displacement (mean ± SD) values for each three receiver cases.

## Discussion

In this study, we developed a biofidelic head-neck FE model from MRI datasets and validated its’ biomechanical responses against four experimental datasets [63–66]. The numerical solutions revealed that our developed model was capable of accurately simulating the experimental impact scenarios and predicted the biomechanical response of model structures (e.g., brain stress-strain tensor values) reasonably well. The results also showed that most geometrical measures were within the normal range of previously-reported values (Tables 4–6). Only scalp thickness and CSF volume of our model were somewhat larger than their corresponding values in the literature [46, 49]. In order to verify that they were not erroneously segmented in the T2 MRI sequence, we also imaged and segmented T-1 weighted images of the same subject and found the same scalp thickness and CSF volume. Not to mention, previous studies reported that skin thickness is correlated with body weight [68]. As our study participant was a 92^nd^ percentile male by weight [45], thus he might have a higher-than-average scalp thickness. Additionally, we defined CSF volume as the space between the skull and brain, which includes dura and pia maters, intracranial blood vessels, meninges, etc. As we did not model other structures except dura and pia maters between the skull and brain, we conjecture that the inclusion of their spaces as CSF led to a larger CSF volume. Six out of 22 cervical vertebral measures (C3 height, C6 and C7 depth, C4 spinous process length, and C3 and C4 vertebral body to spinous process length) [50] and the C5/C6 disc height [51] were found to marginally exceed their normal range, as reported in a previous study [50]. A few previous studies have also reported such geometrical discrepancies in the neck region [15, 40, 50]. Such as, Liang et al. [15] observed larger C2 and C3 spinous process lengths, and Barker et al. [40] found higher posterior disc height than the reported literature data. This leads us to believe that, regardless of the imaging technique and the study subject’s anthropometric distribution, voxel-based intensity standardization methods [69] should be implemented to enhance soft- and hard-tissue contrasts in the complex neck region. The lack of such an image processing method might have led to slight discrepancies in those vertebral measures, even though we took utmost care to segment and reconstruct the 3D geometry of each neck structure.

When a mechanical impact is introduced to the head and neck system, the resultant movements (and/or stress) transfer(s) from one tissue to another and cause deformations in every head-neck structure. Therefore, it is crucial to ensure that our model was able to accurately reproduce these events and assess the mechanical behavior of all head-neck structures. For this purpose, we simulated four experimental studies: 1) NBDL study [63] (high linear acceleration impacts that are commonly seen in automotive and aviation accidents), 2) Nahum’s study (low-velocity projectile impacts [64], that are common in occupational and battlefield-related head traumas) 3) Ito’s study [65] (simulated frontal impact to assess the biomechanical response of the cervical spine), and 4) Alshareef’s sonomicrometry study [66] (brain displacement in response to angular velocity). As we incorporated certain geometric and mechanical simplifications (e.g., linear elastic properties of bones) in our original model, we expected slight differences between in-vivo results [63] and our model predictions. However, the high correlation (r>0.97) between neck flexion angle profiles and the minor difference (5 ms) in their peak time between NBDL experimental and numerical simulations indicate that our model’s head-neck damping characteristics closely approximate those of a living human, which is especially noteworthy since NBDL employed live human subjects rather than cadavers as in other studies (Fig. 4). This highlights the fidelity of our model’s neck damping characteristics to that of living humans.

The numerical results of Nahum’s study also revealed that the disparities between our model’s predictions and the experimental data are relatively minor. For instance, our head-only model’s parietal and occipital intracranial pressure values were very close to Nahum’s experimental (cadaver head) results. Only a slight difference in the frontal intracranial pressure (9% lower) was observed due mainly to the extra thickness of the scalp that absorbed a greater amount of impact energy than usual. Additionally, in the pursuit of validating their head model using Nahum’s experimental ICP data, Sahoo et al. [70] found that the maximum difference in ICP was below 5%. In our study, we observed ICP differences of 3.6%, 2.0%, and 8.1% in the frontal, parietal, and occipital regions, respectively. In comparison to Sahoo et al.’s study results, we observed slightly higher ICP in the occipital region, which may be attributed to the fact that Sahoo et al. incorporated anisotropy properties in their brain model. Despite the high correlation coefficients between the experimental and model-predicted ICP patterns, we observed slight phase difference in the frontal (0.80 ms) and occipital (0.37 ms) regions between our model’s response and the experimental results. This could be due to the fact that the damping properties of brain and/or CSF used in the experimental cadaver (in this case, Nahum’s study) is different from the in-silico modeling of living tissues [71]. Furthermore, the inclusion of neck yielded a greater time lag (phase delay) between the simulated and experimental results, compared to the head-only model. Especially, the neck damping might have contributed in lowering ICP in the frontal and parietal regions. A closer examination of Figure 6 also highlighted that the presence of the neck diminished the dynamic von-Mises stress in the brain, underscoring the crucial role played by the neck in mitigating brain injury. Additionally, the pressure response of the brain, akin to hydrostatic pressure, is notably influenced by its Poisson ratio [72]. The increase in ICP in the occipital region can be attributed to the brain compressibility behavior as we used a Poisson ratio of 0.4977 to model our brain.

For Ito’s study [65], the most significant disparity between our values and the experimental mean was observed for C2-C3 intervertebral disc, especially the posterior region of the disc, where our simulated peak shear strain fell within two standard deviations of the experimental value. Like our study, Barker et al. [41] have also sought to validate their neck response using Ito’s frontal impact experimental results. They observed higher model-predicted peak shear strain values than the experimental results. Particularly, the anterior peak shear strain at C3-C4, C4-C5, and C5-C6 discs and the posterior peak shear strain at C3-C4 and C4-C5 discs were higher than one standard deviation. This variation in the peak shear strain can be attributed to the use of an artificial head in the experiment. Although our model’s head possesses similar weight and moment of inertia characteristics to the artificial head used by Ito et al., the proximity of the C2-C3 disc to the head could magnify the impact of the artificial head’s influence on the results. Furthermore, the Ito’s experimental study used constant and equal force springs for muscle activation, differing from our approach of uniform and constant activation levels, potentially contributing to slight variations in the peak strain values across all intervertebral discs.

While comparing the brain displacement results of our model with Alshareef’s experimental values [66], we found a high correlation in the Y (medio-lateral) and Z (superior-inferior) directions and a medium level of correlation in the X (anterior-posterior) direction (Fig. 8). As Alshareef’s experimental motion occurs in the Y-Z plane (lateral direction), the displacement in the X-direction should ideally be zero in all cases. However, both experimental and simulation patterns of the brain displacement kinematics showed trivial amount of fluctuation (non-zero values) in the X direction, which contributed to a comparatively lower correlation values in the X direction. Additionally, the absolute brain displacement values in the Y direction for Receiver 9 was also found to lie outside one standard deviation of the experimental value. Previous FE study by the same author has also found such discrepancies, where they reported a CORrelation and Analysis objective rating system (CORA) score of 40-60 between their numerical results and Alshareef’s experimental data [37]. Furthermore, as the brain displacement kinematics depends on the brain-fluid-skull relative motion, a realistic modeling of brain-CSF and CSF-skull interfaces is crucial to obtain accurate brain displacement results. However, modeling the CSF as a fluid and the CSF-skull and CSF-brain interfaces as complex fluid-structure interactions with a no-slip boundary condition are challenging [73]. Thence, previous studies had modeled CSF as a linear or nonlinear solid material with low stiffness, high bulk modulus, and high Poisson’s ratio to mimic its’ fluid-like behavior [9] and its’ interactions with other tissues as tied or sliding contacts [9, 10]. When the CSF encounters adjacent stationary tissues in the brain, it comes to a complete stop (zero velocity) at the contact surfaces. This closely resembles a tied contact approach in solid mechanics. As a result, the brain displacement kinematics (i.e., the brain-skull relative motion) was primarily the deformation of the CSF since we implemented a tied contact between CSF-brain and CSF-skull tissues.

This study had several limitations. First, we segmented our brain model into gray and white matter but assumed isotropic properties for both, neglecting anisotropy in brain tissue. Second, we simplified the model by considering identical, constant muscle activation for extension and flexor muscles which are different from brain-controlled actual muscle activation patterns. Third, we did not implement voxel-based intensity standardization in the image processing, leading to slight discrepancies in 3-D vertebral body geometry. Fourth, both skull and vertebral bodies were not separated into cancellous and cortical bones, and the CSF was not modeled as a pure fluid. Future works may make the model more biologically accurate by modeling cortical and cancellous bones separately and CSF as a fluid and CSF-skull and CSF-brain interfaces as fluid-structure interactions. Fifth, we used four widely recognized experimental studies for validating our model’s head and neck response. However, it’s worth noting that there are other available experimental studies, especially experimental studies on sensor-based brain strain measurement. These studies can be used for further validating the robustness of our model against various other types of head impacts. Sixth, in this study, we did not scale the model to maintain the geometric and structural integrity of the study subject. As the mechanical response of the brain depends on its size, some discrepancies between the model response and experimental data (Alshareef’s and Nahum’s studies) can be attributed to the differences in experimental and modeled brain size. Despite these limitations, our developed head-neck model was able to reproduce the experimental results mostly and provide valuable insights into the mechanical responses of the brain both with and without the presence of neck structures.

In summary, though there exists numerous validated head-neck model, we believe that this MRI-based computational FE platform composed of detailed, biofidelic head (includes scalp, skull, pia and dura maters, CSF, brain’s white and grey matters) and neck (consists of C1-C7 cervical spines, discs, 14 ligaments, and 42 neck muscles) structures, an innovative hexahedral meshing technique for the brain and CSF, and a scalp erosion model make notable contributions to advancing the computational injury biomechanics research and has the potential to be used as a computational tool for various applications, such as brain injury prediction, helmet design, motor vehicle safety, and ballistic head impact injury predictions.

## Acknowledgments

This work was partly supported by the U.S. Department of Homeland Security (70RSAT21CB0000023). The MRI data acquisition was supported by the Texas Tech University Neuroimaging Center.

